# Noise-driven cell differentiation and the emergence of spatiotemporal patterns

**DOI:** 10.1101/220525

**Authors:** Hadiseh Safdari, Ata Kalirad, Cristian Picioreanu, Rouzbeh Tusserkani, Bahram Goliaei, Mehdi Sadeghi

## Abstract

One of the major transitions in evolution is the step from unicellularity into the brave new world of multicellularity. To understand this feat, one has to fathom two main characteristics of multicellular organisms: differentiation and self-organization. Any explanation concerning this major transition should involve mechanisms that can simultaneously explain the marvellous intricacies manifest in the aforementioned characteristics, and an account of the evolution of such traits. Here we propose a noise-driven differentiation (NDD) model. The reliance on noise, in place of a more mechanistic approach, makes the NDD model a more suitable approach to explain differentiation and self-organization. Furthermore, our model sheds some light on the possible evolutionary origins of these biological innovations. To test the NDD model, we utilize a model of cell aggregation. The behavior of this model of cell aggregation is in concert with the NDD model.

## Introduction

The traditional idea of a living cell where every organelle, every reaction, and every interaction is part of a clock-like order has long been shattered by the understanding that biological systems usually struggle to function in noisy environments. One might consider life to be an uphill battle against pandemonium, where disarray is the norm and spheres of order – i.e., biological systems – are rarities that are unlikely to appear in the first place. In this view, noise is a nuisance that natural selection always attempts to eliminate. It is for the same reason that selection cannot increase the fidelity of replication beyond a certain threshold; the biological cost of increasing fidelity simply becomes too high at that point (***Kimura, 1967***).

A different view has recently gained some grounds (***Balázsi et al., 2011***; ***Chalancon et al., 2012***; ***Huang, 2009***; ***Losick and Desplan, 2008***). In this view, biological systems that regulate and utilize the noise can have higher fitness under certain circumstances. Had biological systems been utterly deterministic, adaptation – i.e., the emergence of a new phenotype or a change in the gene expression pattern to utilize a new food source – would have been impossible without the emergence of new mutations. In reality, noise in the cell can result in beneficial non-genetic diversity in otherwise genetically homogenous populations – e.g., cyanobacteria (***Wolk, 1996***) and yeast (***Paliwal et al., 2007***). But what mechanism can account for the presence of phenotypic diversity amongst daughter cells that are genetic clones of each other? Is it possible for a stochastic mechanism to explain the non-genetic diversity? Even if such stochastic explanation were offered, how could this explanation possibly account for the ordered spatiotemporal patterns in spatially-extended cell population?

The model of cell differentiation proposed in this work, henceforth referred to as the noise-driven differentiation (NDD) model, accounts for the peculiarities of this biological phenomenon by weaving noise into an explanation of cellular behaviors at the time of differentiation. While on the surface, this approach might seem lofty and even radical, the model discussed in this paper is parsimonious when it comes to the mechanisms requisite for its operation. The NDD model rests on 8 components(Table 1). Some can be regarded as facts, based on reliable empirical evidence from biological systems (components #1 #2), while others are accurately described as assumptions (components #3 – 8).

**Table 1.** The components of the NDD model.

| # | Component | Justification |
| --- | --- | --- |
| 1 | Noise, resulting from a plentitude of sources, is an inseparable part of a living cell. | Based on the observed effect of noise on the processes in living cells, from microbes to mice (e.g., see *(Kepler and Elston, 2001; Ozbudak et al., 2002; Elowitz et al., 2002; Maamar et al., 2007; Chang et al., 2008)*). |
| 2 | Stochastic partitioning of cytoplasm during cell division and the random distribution of molecules in the cytoplasm determine the cytoplasmic contents of the daughter cells. | Variation in the position of cell-division plane is a biological fact (reviewed in *Margolin (2000); Monahan et al. (2014); Pickett-Heaps et al. (1999); Wu and Tzanakakis (2012); Bradshaw and Losick (2015)*), and its effect on the diversification of cells is well-known (e.g., see *Jan and Jan (1998); Betschinger and Knoblich (2017)*). |
| 3 | The fate of a cell is determined when it is born. | Based on the assumption that cell-fate-determining factors are in small numbers in a cell and the stochastic distribution of these factors during cell division determines the fate of the newly-born daughter cells. |
| 4 | Cell fate is determined by a switch. | Genetic switches have been observed in a variety of taxa (reviewed in *Balázsi et al. (2011)*), and has been proposed as a model to account for cell differentiation (e.g., see *Perez-Carrasco et al. (2016)*). |
| 5 | The interaction between the building blocks of the switch determines its bias. | Our assumption based on our knowledge of well-known genetic switches, such as $\lambda$ phage (see *Cortes et al. (2018)*). |
| 6 | All the information needed to construct the switch is genetic. | We assume that, while stochasticity is what drives the decision made by the switch, the information necessary to construct the switch is encoded in the genetic content of a cell. |
| 7 | The robustness of the switch is the result of a complex network of interactions. | Our assumption based on *Sharifi-Zarchi et al. (2015)*. |
| 8 | Cell fate is influenced by its location and its environment.* | We assume the the switch determining cell fate should, in addition to being swayed by the intrinsic factors, be influenced by its neighbors. |

\* This component is necessary for the ordered spatiotemporal patterns in cell population.

There is a plethora of phenomena within a cell that can contribute to its intrinsic noise – e.g., transcription regulation, transcription factor binding to the DNA, RNA processing in eukaryotes, translation, post-translational modifications, protein complex formation, protein and RNA degradation, etc. Single-cell level measurements of gene expression further cements the notion that cells are intrinsically noisy when it comes translating its genotype into phenotype (***Sanchez and Golding, 2013***). The displacement of the division plane relative to the middle of the cell can result in an unequal distribution of cell content between the daughter cells, even if molecules are homogeneously distributed within the cell. In fact, the central role of asymmetric cell division in the diversification of cells, from *Drosophila* to mammals has been known for many years (***Jan and Jan, 1998***; ***Betschinger and Knoblich, 2017***). The components #1 – 2 is an acknowledgement of the role stochasticity in living systems based on these observation.

Thus far, two types of solutions to the problem of cell differentiation have been proposed: the first category consists of models that rely on cell-cell communication (reviewed in ***Wolpert*** (***2011***)) and the second category relies on asymmetric cell division (reviewed in (***Rudel and Sommer, 2003***)). The research project within the confines of the former category is mainly a quest to find the building blocks of the apparatus that makes the specific kind of cell-cell communication needed for cell differentiation. The latter category, on the other hand, presumes the asymmetric cell division to result in differentiation. Hitherto unknown and often complicated mechanisms have been proposed to explain the asymmetric distribution of fate-determining factors during cell division (***Morrison and Kimble, 2006***; ***Clevers, 2005***). Both categories are quintessentially mechanistic in nature, since they rely on mechanical interactions at the cellular level. While we agree with the importance of the asymmetric cell division, it seems to us that a stochastic model of differentiation, like the NDD model, negates the need for new mechanisms. In this model, we adopt the view that stochastic processes result in differentiated cells due to the distribution of key proteins, instead of cells differentiating by receiving signals after they are born (component #3).

The component #4 is based on the idea that characteristics of a cell can be changed by a switch (Not a very recent idea, e.g., ***Novick and Weiner*** (***1957***)). The notion that cell fate is determined by a switch is best illustrated by the now famous case of the *λ* phage. The process by which the phage decides to integrate into the host’s genome – i.e., lysogenic – or to replicate copies of itself in the cell until it bursts open – i.e., lytic – can be explained by a stochastic switch which makes that portentous decision in a probabilistic fashion, while taking into account the presence of certain key factors (***Ptashne, 2004***). One can assume that the bias of this switch is determined by the interactions of its building blocks (component #5). For example, upon infecting bacterial cells, *λ* phage proceeds to lyse the host, but as the concentration of CII protein increases, so does the likelihood of the reactions suppressing the activation of *pR* and *pL* promoters, relevant to the onset of the lytic trajectory, which in turn, tilts the scale away from lysis towards lysogeny (***Cortes et al., 2018***). We propose that phenotypic diversity arises from the effect of the noise on a genetic circuit that exhibits a switch-like behavior (component #6). The notion that different phenotypes are produced from the same genotype as a consequence of noise is widely observed in nature (reviewed in (***Vogt, 2015***))

How robust can a fate-determining toggle switch in the face of new mutations? ***Sharifi-Zarchi et al.*** (***2015***) took advantage of the gene expression profiles of **442** mouse embryonic cells to construct a network of key transcription factors (TFs). While a regulatory circuit with two TFs could explain differentiation, They reasoned that such a simple switch is susceptible to mutations. To construct a robust switch, they built a circuit with two clusters of TFs with correlated expressions. Expectedly, the alternative switch, which involved more interactions, was much more robust. We would expect different levels of robustness for a switch, given its biological importance in evolution (component #7).

The components #1-7 are sufficient to generate a population of cells with different proportions of two phenotypes (Fig 1). While this kind of fate determination is adequate vis-à-vis primitive cells with no organization, it does not allow the emergence of multicellularity. An additional component is necessary to explain this major transition from mere phenotypic differentiation to ordered spatiotemporal patterns in the body of a multicellular organism. For self-organization to occur, we assume that the toggle switch determining cell fate should, in addition to being swayed by the intrinsic factors, be influenced by its neighbors (component #8).

**Figure 1.**
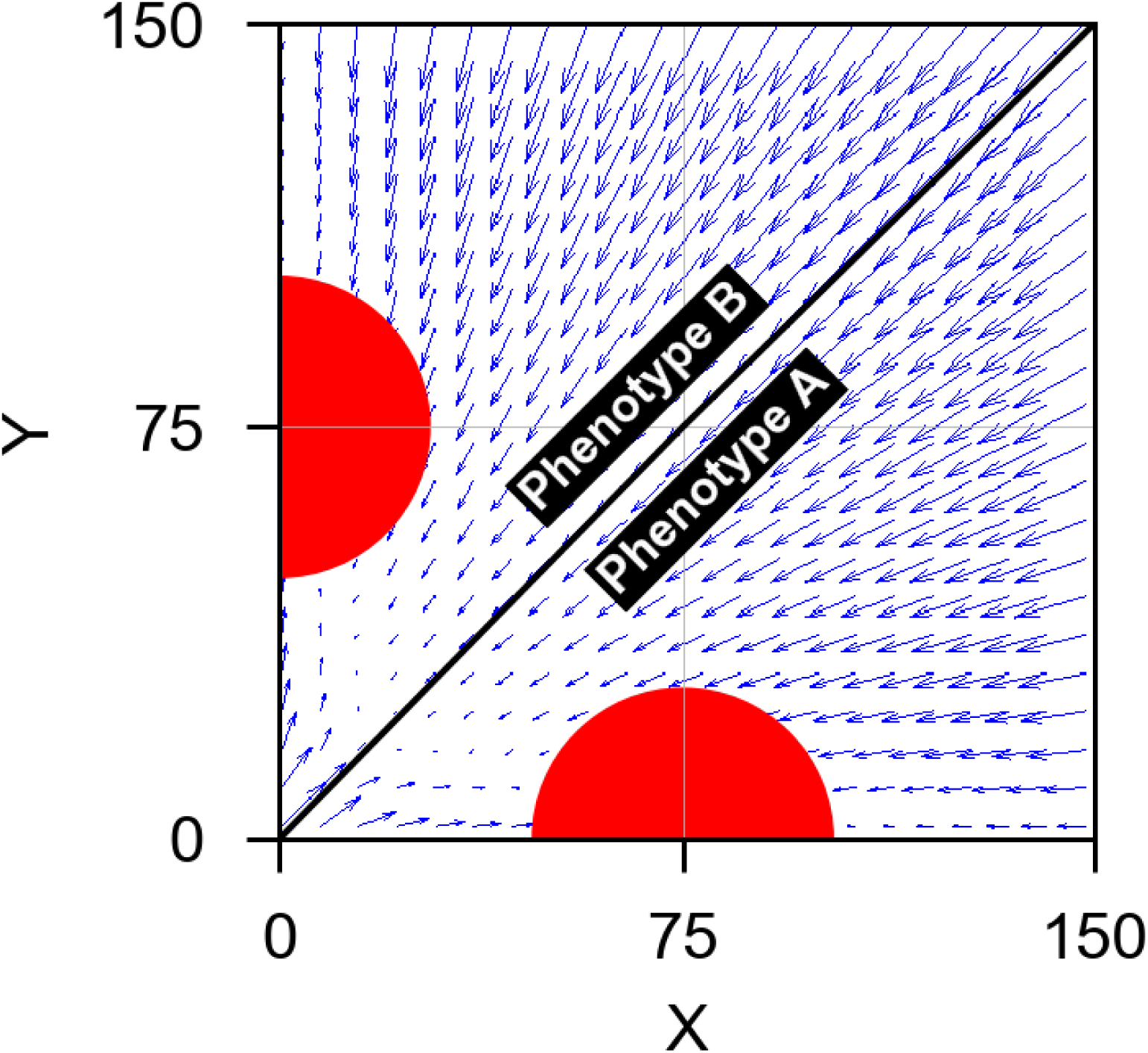
The phase-portrait diagram for the NDD model (based on Eq 1). In a bistable switch, two attractors (red semicircles) and, consequently, two phenotypes are available:***A*** and ***B***. The likelihood of a switch choosing state ***A*** over ***B*** depends on the number of the transcription factor associated with state ***A***(**TF** _*X*_) relative to the number of the transcription factor associated with state ***B***(**TF**_*Y*_), as well as the noise in its environment. The parameters used to generate this and the following figure are as follows: *n*=2, *β* = 0.1, protein half-life = 10min, and protein dissociation constant = 10. Unless noted otherwise, these parameters are used in all the subsequent figures.

To test the general veracity of the NDD model, we used a simple model of cell aggregation. In this model, a simple switch is defined that can switch between phenotypes, ***A*** and ***B***.

## Results

The overall behavior of the cell aggregation model demonstrates the principles of our framework – that is, the stochasticity results in phenotypic heterogeneity as the population grows in size (movie S1). To further illustrate how each source of noise affects the cell differentiation, we focused on each source separately in the simulations.

### The stochastic positioning of division plane and the stochastic distribution of key proteins affect differentiation

One source of intrinsic stochasticity stems from the random positioning of the division plane. This factor would disproportionately influence the number of molecules that exist in low numbers within cytoplasm. In this work, it has been postulated that the determinants of cell fate are low in numbers and thus, greatly affected by stochasticity.

To demonstrate this phenomenon, the position of the division plane was allowed to vary with respect to the mid plane of the cell. Starting from a cell with phenotype ***A,*** in which the protein ***X*** is dominant, the population heterogeneity –i.e.,emergence of phenotype ***B*** – was traced over 12 generations. The results are shown in Fig 2. When the division plane is situated in the middle of the cell, and the TFs are relatively abundant, very few cells differentiate. As the variance in the cell-division plane increases, so does the proportion of ***B*** cells. This phenomenon is dependent on the number of proteins, since such bias is more pronounced when the number of proteins is relatively low. In fact, with large numbers of TFs in a cell, it will be more likely for its daughters to have almost the same density of TFs as their mother. Thus, they will be in the same domain as the mother in the phase space, and their fates will be identical to hers. This can be seen clearly in the lower curves in Fig 2. However, for low copy numbers of TFs, the difference between TF numbers in two daughter cells becomes more prominent and can even lead to different cell fates. Therefore, it is possible to have heterogeneity in the population in the absence of any other noise, i.e., cells with low TF numbers are heterogeneous even with no variance in division-plane displacement (Fig 2a). Adding spatial fluctuation to the distribution of TFs within a cell increases the chance of differentiation, since in this case, in addition to the noise from the positioning of division plane, the key proteins are stochastically distributed as well (Fig 2b).

**Figure 2.**
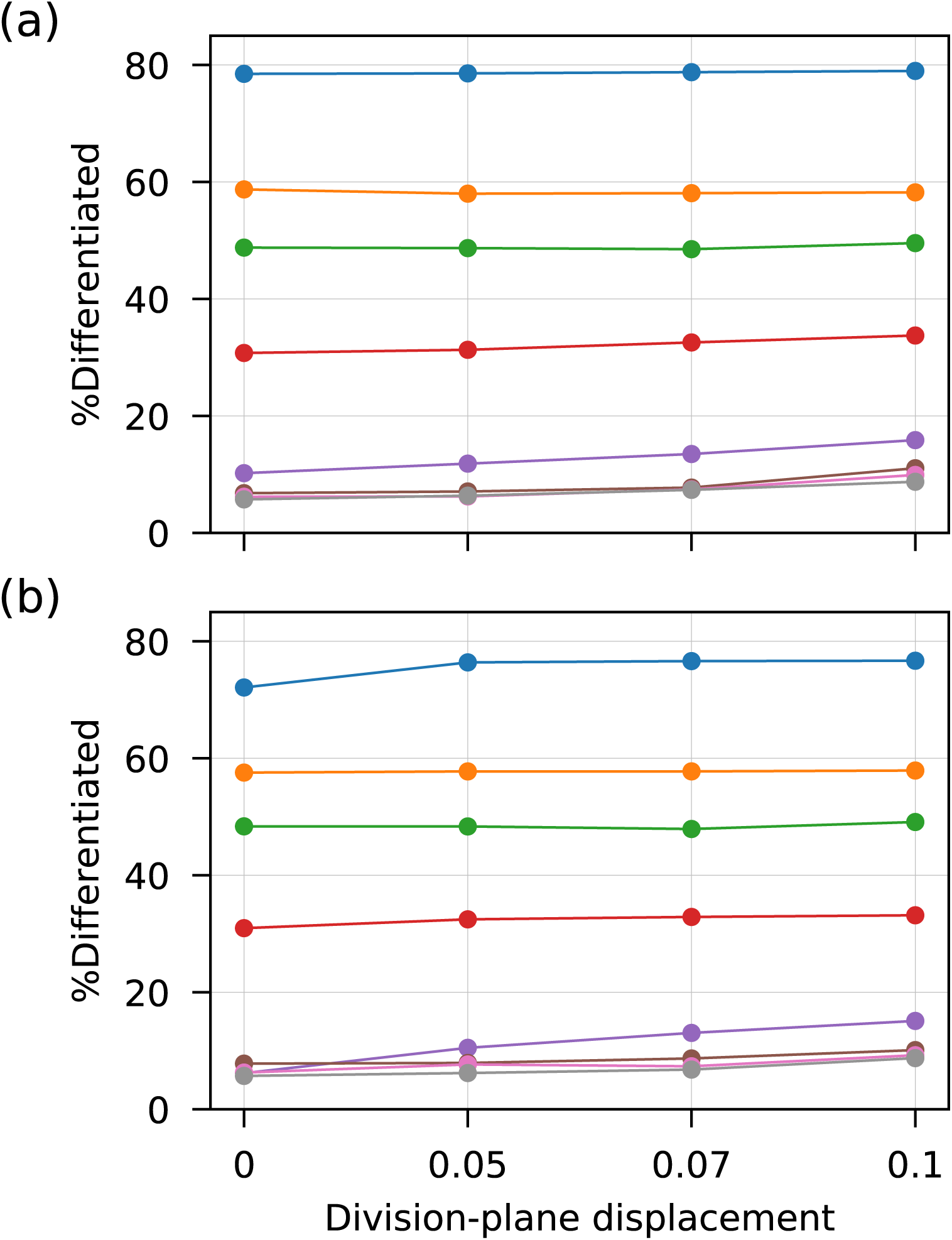
The stochastic positioning of the division plane and the random distribution of TFs in the cytoplasm, as intrinsic sources of noise, affect the none-genetic phenotypic diversity (component #2). The phenotypic diversity is represented by the proportion of cells with the phenotype ***B*** relative to the total number of cells in the population. In panel (a), the only source of noise is the stochastic positioning of the division plane, while panel (b) shows the phenotypic diversity as a result of both sources of noise. In each panel, the curves indicate different amounts of protein *X* in the mother cell; from top to bottom, respectively, *X* = 10, 15, 20, 25, 35, 45, 55, 100. The results are average over 100 replications. Error bars are 95%Cl

### Signaling can create spatial order

In the cell aggregation model, ***B*** cell can release signals in the environment. These signals diffuse at a slow rate and, consequently, have a very short radius of influence. The absorption of these signals by other cells in the population affects the number of proteins involved in the switch – that is, switching to the phenotype ***B*** during cell division becomes more likely (Fig 3). When this environmental signaling is added to the population, the cells organize in a non-random fashion, a stark contrast to the random heterogeneity observed before (Movie S2).

**Figure 3.**
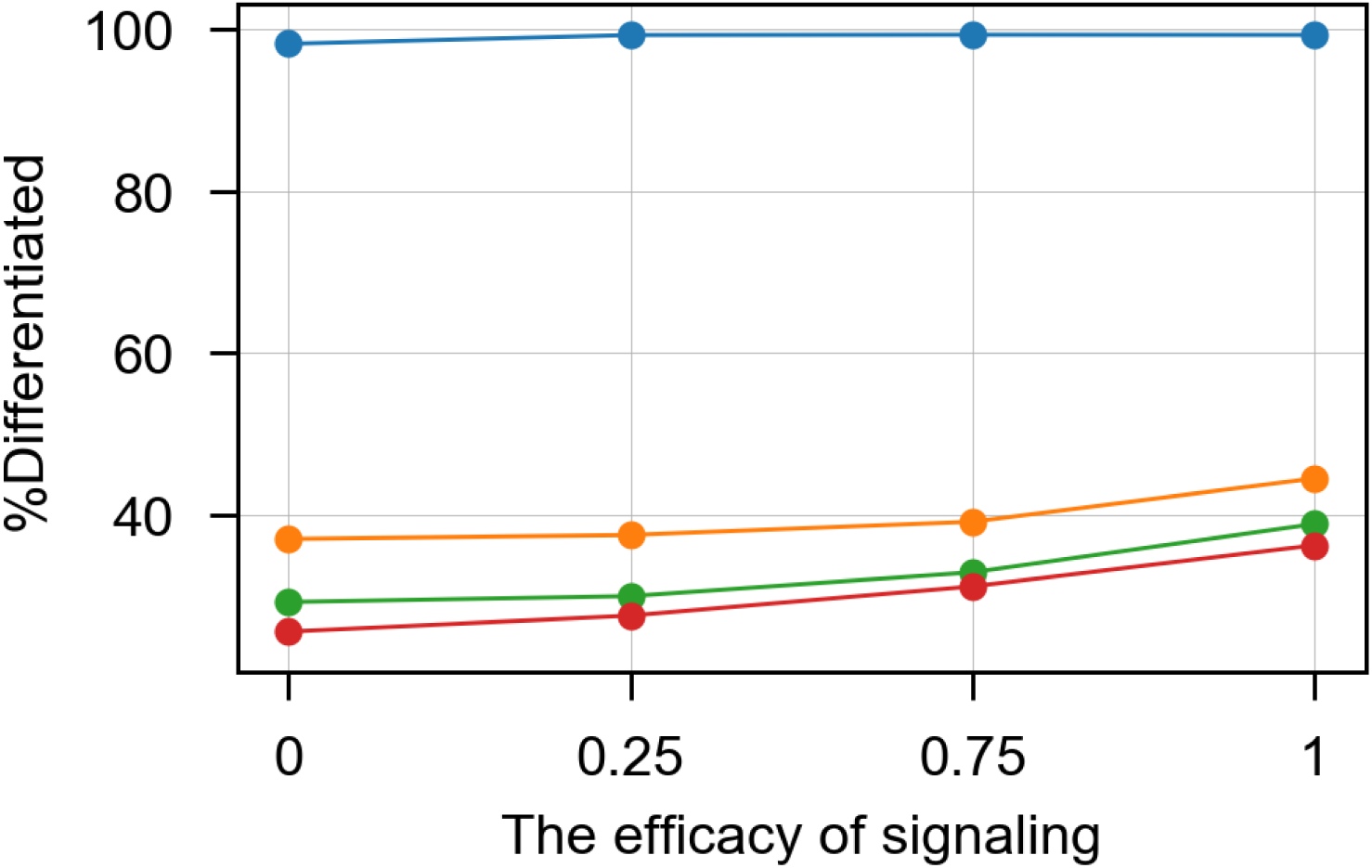
Adding signaling to the cell aggregation model results in higher none-genetic phenotypic diversity, compared to populations without signaling (as shown in Fig 4). The phenotypic diversity is represented by the proportion of cells with the phenotype *B* relative to the total number of cells in the population. The curves indicate different amounts of protein in the mother cell; from top to bottom, respectively, *X* = 10,35,55,100. It fascinating to notice how the lowest number of TFs (*X* = 10) results in total differentiation. The efficacy of signaling is defined as follows: if in the position of a cell with phenotype ***A***, the signal concentration exceeds the mean signal concentration, then this cell would have more chance of becoming a ***B*** cell. The results are average over 100 replications. Error bars are 95%CI.

Fig 4 represents a visual understanding of the results from the NDD model. It shows the bacterial community in a 2-dimensional simulation area after more than 8 generations. In Fig 4a, the variance in the stochastic positioning of the division plane increases from left to right. It can be seen that the heterogeneity in the population increases as well by the presence of new phenotypes (cells in orange). In Fig 4b, development of an organized community as a result of signaling molecules is apparent (group of orange cells). The organization observed will increase over time and the community of orange cells will develop (Movie S3).

**Figure 4.**
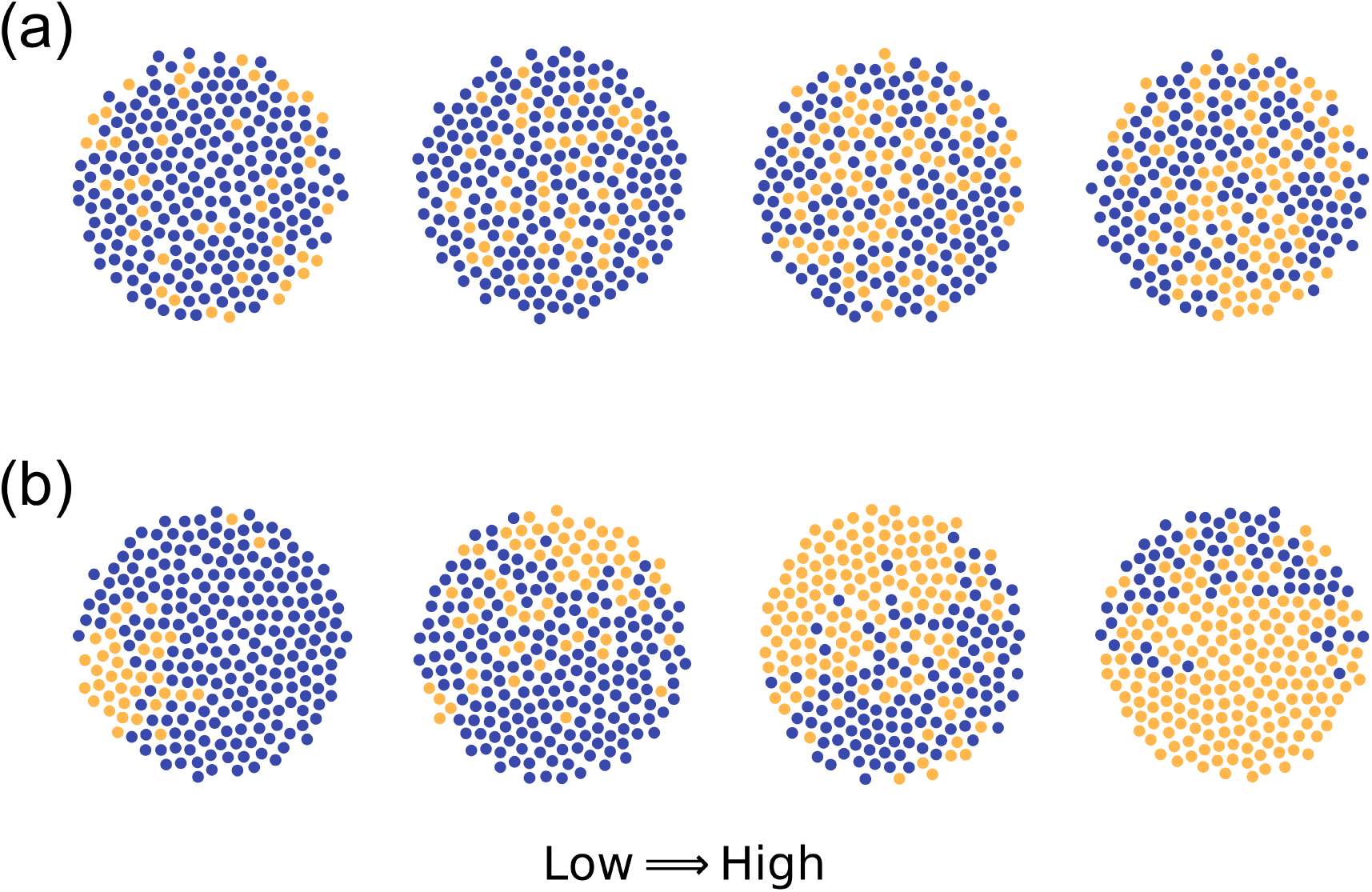
Population heterogeneity as a result of the noise in: (a) the number of TFs in the daughter cells or (b) the secretion of signals from the ***B*** cells. Both the number of TFs in (a) and the efficacy of singling in (b) increases from left to right in this figure. The blue circles represent the ***A*** cells and the orange ones represent the ***B*** cells. Each aggregation is the final state of a single run of the stochastic model with the given parameters. The amount of protein *X* in the initial cell in each simulation was 35. The radius of the area of aggregation is 100*μm.*

## Discussion

Molecular processes in the cell are noisy events that result in varying degrees of heterogeneity. Taming this inherent noise is vital for the emergence and the continuation of life. In fact, life can be characterized as a system with the capacity to control noise. The phenotype of a cell is generally stable, but during cell division, this cell can produce daughter cells with different phenotypes via symmetric or asymmetric cell division. The resulting non-genetic phenotypic diversity is a way to achieve adaptation in a fluctuating environment by producing phenotypically diverse offspring without any need for genetic change. Given the variety of sources of noise, the cell fate determination can be a stochastic process. One can imagine a few genes involved in cell fate determination, where the noise in the cell affects the proportion of daughter cells born with a certain phenotype. The ability to change the phenotypic proportion of daughter cells via a stochastic mechanism, which is also tunable, is a superb strategy to outcompete rivals bereft of such gift.

Given the prominence of noise in living cell, we argue that the NDD model can provide a satisfactory explanation of how organization can emerge from noise. Proposing a stochastic model of cell differentiation is not an entirely novel concept, e.g., see ***Suzuki et al.*** (***2011***); ***Yamagishi et al.*** (***2016***) as examples of an impressive body of work produced by Kunihiko Kaneko and his colleagues on this subject and ***Kupiec*** (***1997***); ***Paldi*** (***2003***) as similar proposals regarding the possible role of stochasticity in generating phenotypic diversity. We argue that our approach differs from theirs and similar ideas in certain important aspects: firstly, our model assumes that cell fate is determined when the cell is born, and secondly, that stochastic fluctuations in the cell, and the effect of signals from neighboring cells in the multicellular case, drive the phenotype of the cell towards one attractor rather than another during cell division. This approach is in keeping with the recent emphasis on the importance and the prevalence of noise in biological functions, specifically cell fate (***Balázsi et al., 2011***; ***Huang, 2009***; ***Kittisopikul and Süel, 2010***). The model of cell aggregation used in this study allowed us to test all the components of the NDD model, barring components #6 and #7, which demand through investigations of their own. This model of cell aggregation provides us with a relatively realistic depiction of the process that results in phenotypic differentiation in a population. We believe that, with few changes, the NDD model can be applied to other biological systems as well.

The ability of the cells to differentiate into different types was the crucial step that enabled the ancient solitary cells to leave the primordial soup behind and evolve into the vast array of specialized cells we see today. As ***Queller and Strassmann*** (***2009***) point out, there are different shades of organismality –i.e., the ability for components to work together with little conflict among them–, each shade resulting from the affinity of the members of the system to cooperate versus the temptation to cheat. We can sidestep the problem of conflict since in prokaryotic multicellularity, e.g., biofilm, and in most truly multicellular eukaryotes, the cells are highly related, thus lowering the probability of cheating (***Ostrowski and Shaulsky, 2009***). Without tangible levels of conflict, multicellularity as a trait becomes patently advantageous. In their seminal work, Maynard Smith and Szathmáry (***Maynard Smith and Szathmáry, 1995***) considered two possible mechanisms to account for the emergence of cell differentiation: one relies on the presence of determinants that prohibit the stem cell to differentiate, and the other postulates the cell-cell contact as a mechanism that determines cell fate. While these suggestions account for how the multicellularity might be sustained, they do not explain how this major evolutionary transition could have occurred in the first place.

It is easier for cell differentiation to evolve via the emergence of a switch, rather than the less plausible path that involves the evolution of a clockwork mechanism. According to the NDD model, the emergence of early stages of multicellularity only requires the evolution of a suitable switch – the rest of the necessary ingredients needed for the transition into self-organization is provided by the stochastic elements affecting the switch. The major transition from unicellularity to multicellularity –i.e., from phenotypic diversity in a population to from an ordered and stable spatial heterogeneity– only requires one more step: the evolved switch should be simply affected by the signal(s) released by its neighbors (components). The spatial information received in this way would bias the switch such that the population-level organization is retained. It is tempting to postulate a connection between the cell-differentiation switch, postulated in the NDD model, and the toggle switch used in quorum sensing in bacteria (***Hooshangi and Bentley, 2011***). Quorum sensing enables bacteria to regulate their phenotypes apropos of their neighbors and is more robust in a dense community (***Schluter et al., 2016***). It seems plausible to consider this type community-based phenotypic regulation as a precursor to similar switch-based mechanisms for cell differentiation in multicellular organisms.

In their criticism of a noise-driven alternative to their model, ***Suzuki et al.*** (***2011***) considered it unlikely for a noise-driven model to maintain the exact levels of stochasticity needed to produce the desired proportion of differentiated cells to stem cells. In our view, this conclusion follows from a non-evolutionary perspective, since it is easy to imagine negative selection keeping a genetic switch just sensitive enough to result in a correct differentiation pattern vis-á-vis the biological fitness. Furthermore, if a switch is robust (component #7), then it will be able maintain its bias in the face of new mutations. ***Suzuki et al.*** (***2011***) also point out that a noise-driven model can only produce reversible differentiation. While the NDD model as described here only explains the phenotypic differentiation in prokaryotes, which is indeed reversible, it seems that changing the bi-stable switch to a tri-stable one could remedy this issue and explain the irreversibility of differentiation observed in eukaryotes, as it should increase the strength of attractors (***Ghaffarizadeh et al., 2014***).

One of the quintessential aspects of the discussed model is its population-level perspective. Population-level thinking is one of the main points of the evolutionary theory, and bringing it to explain a cellular phenomenon can lead us to reap valuable insights. While a population of cells has, on average, certain properties relevant to differentiation, e.g., the mean number of key proteins, the average position of cell division plane, and etc., these average values do not tell the whole story. Instead, the variance in these values, i.e., the non-genetic variation present amongst individuals, is the key to understand differentiation (as observed in studies such as in ***Chang et al.*** (***2008***); ***Moussy et al.*** (***2017***)). This noise in the population is essentially the fuel that propels cellular differentiation, be it in the reversible differentiation in prokaryotes or the more complicated irreversible ones in higher organisms. We believe that this population-level vintage point is the necessary tool to understand this otherwise mind-boggling biological process. Without this perspective, the task of explaining such a seemingly fine-tuned process devolves into an attempt to come up with complex cellular interactions that would make climbing this improbable biological mountain feasible.

The NDD model can be used wherever there is cell division and differentiation. The differentiating cell can be a prokaryotic one, able to divide into daughter cells with dissimilar, and reversible, phenotypes or a eukaryotic cell undergoing irreversible differentiation, without the need for one or a few complicated mechanisms. The transition from single cells into the brave new world of multicellular entities could have been the result of a mechanism very much akin to the NDD model. Such transition is possible because the bias of the switch can be affected by the neighboring cells. The NDD model paints a simple and elegant picture of differentiation and organization, from prokaryotes to eukaryotes. Our model is the logical extension of earlier ideas describing the role of stochasticity in phenotypic variation and the switch-like behavior of genetic circuits vis-à-vis differentiation and multicellularity (e.g., see ***Nanjundiah*** (***2016***)).

## Materials and methods

In the cell aggregation model, the population is made up of cells, where each cell is a circular particle defined by its state variables – e.g., spatial position, size, and phenotype. The simulation geometry is a *L* × *L* square and no flux boundaries. It is assumed that the relative amount of two key transcription factors, *X* and *Y*, controls the cell types; hence, in this model, a cell can have two phenotypes, *A* and *B*, as shown in Fig 1. The dominance of protein *X* leads to phenotype and the dominance of protein *Y* results in phenotype *B*. In fact, a positive feedback loop influences the decision-making process. Two negatively coupled repressors mutually inhibit the expression of the gene that encodes the other repressor-i.e., a toggle switch (component). The rate of this mutual repression is represented in the form of a Hill function (***Gardner et al., 2000***). This positive feedback loop results in two stable steady states, hence implies non-linear approaches. Nonlinear differential equations govern the changes in the number of the repressor proteins, *X* and *Y* (Fig 1);

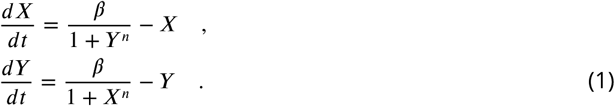

Here, *β* is the effective rate of protein synthesis and *n* is the Hill coefficient, which represents the degree of competence. The number of repressors are represented in the unit of their dissociation constants and time is rescaled by degradation rate of proteins (***Gardner et al., 2000***; ***Carson and Cobelli, 2000***; ***Elowitz and Stanislas, 2000***). Biologically-reasonable values were chosen for the parameters used in our simulation such that Eq 1 would be bi-stable (following (***Gardner et al., 2000***)). This bistable regulatory network has two attractors corresponding to its stable steady states. Based on the amount of proteins at the cell division time, the cell can be in the domain of each attractors, which determines its fate. Depending on the intensity of inhibitory effects of TFs (through the values of constants in the Hill function (***Gardner et al., 2000***)), the two domains of attractors could be equal or not (component #5). Fig 5 shows an example of such behavior in our cell aggregation model. Movie S1 shows the changes in the distribution of TFs in cells around their attractors during the emergence of generation 12.

**Figure 5.**
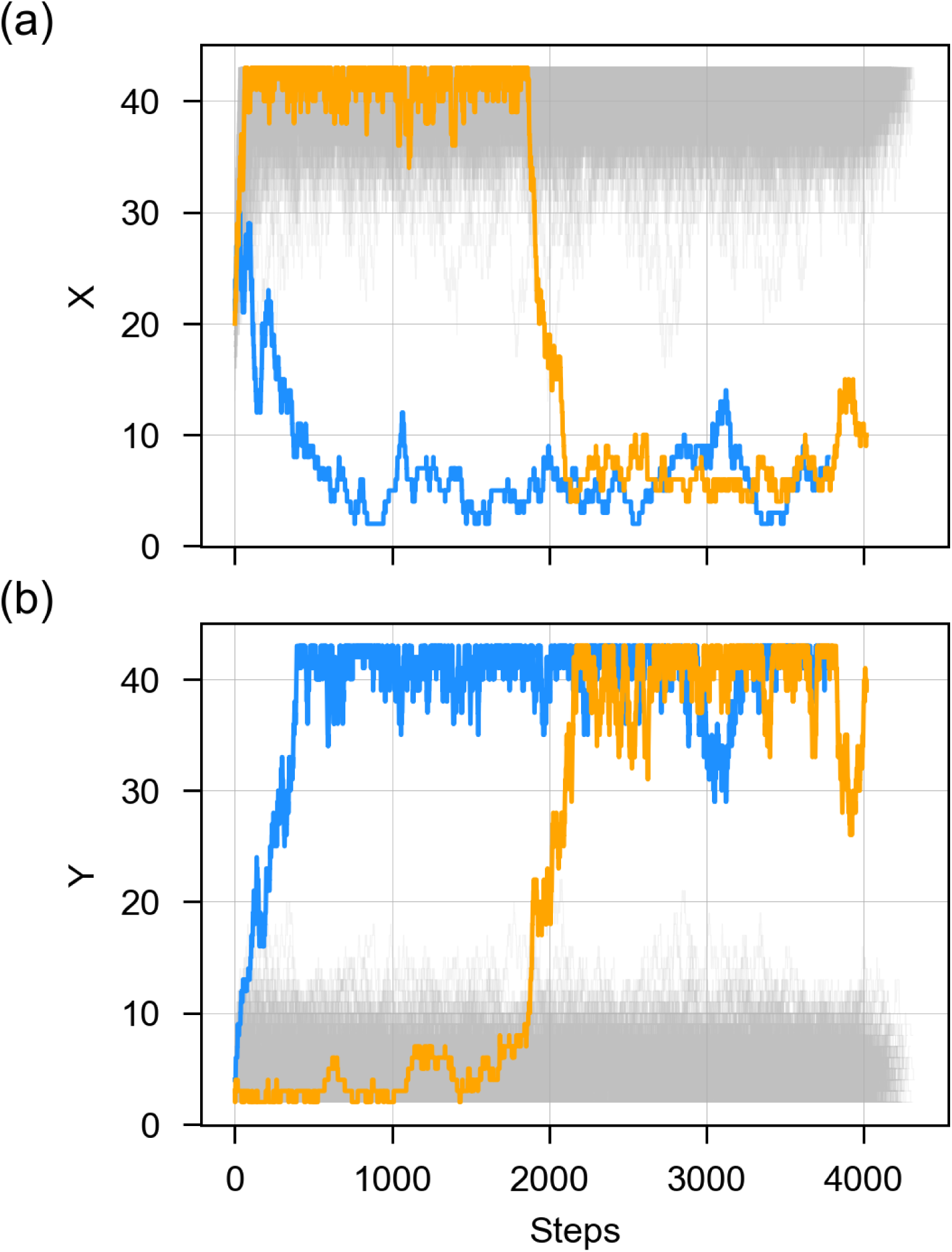
As cells grow, they stochastically explore the phase-plane around their attractor (as depicted in Fig 1) – i.e., over time the values for transcription factors ***X*** (a) and ***Y*** (b) for each cell fluctuate around the attractor that was determined when the cell was born. These fluctuations can result in a cell moving away from its original attractor towards the other attractor, such that it will be more likely for its daughters to have phenotypes different from their parent (the blue and orange trajectories). Results are based on 512 cells that descended from a single cell in the cell aggregation model. The trajectories follow the TFs counts during their lifespan.

## Population growth algorithm

Simulation starts with a single cell with phenotype A. Each iteration in the simulation can be divided into four steps:

1. *Cell growth*: In this step, cells grow linearly in size. Simultaneously, the cytoplasmic content of each cell fluctuates in a stochastic fashion (component #1). The repressor proteins inside the cytoplasm interact with each other and their numbers, ***X*** and ***Y***, are updated; however, because of their low copy numbers, instead of deterministic equations (Eq 1), their fluctuations are captured by the Gillespie algorithm (***Gillespie, 1977***) as a stochastic dynamics for discrete values. According to this algorithm, a probability of occurrence will be assigned to every biochemical reaction in the system. Every protein (*X* or *Y*) is produced with a probability according to the first term on the right hand sides of the Eq 1. As the number of protein ***X*** increases, it further represses the production of protein *Y* and vice versa. Every protein degrades according to its number. In every step of the Gillespie algorithm, one of the above reactions occurs and the time will be updated. The process continues until the number of proteins reaches a steady state.
2. *Cell division*: Even after the number of proteins in a cell reaches the steady state, the cell continues to grow. The growth stops only after the cell reaches a critical size. At this point the cell divides into two daughter cells. The content of the mother cell is distributed among her daughters according to a uniform distribution. In reality and in the presence of active transportation, one can still expect a uniform distribution of molecules in the cytoplasm (***Huh and Paulsson***, ***2011***), making this assumption biologically reasonable. The position at which cell division occurs is randomly chosen based on a normal distribution (component #2). At the time of birth, the phenotype of each newborn cell is determined based on the cytoplasmic contents (number of key proteins, *X*, and *Y* at the time of birth) inherited from the mother cell (component #3). During the cell growth, the number of each protein has a stochastic trajectory in the domain of its attractor and finally it will reach its steady state. In this model, phenotypic change is reversible, meaning that the phenotype can change between the two possible states over generations. Since in our simulations, daughter cells have similar volumes, we consider the number of proteins distributed between them, and not their concentrations.
3. *Relaxation*: After a cell divides, the cells push each other outwards to make room for the new daughter cells (***Kreft et al., 2001***). Simulation proceeds by repeating the steps #1-3. It is worth noting that, without considering self-organization, the process described above would result in a disordered blob of cells.
4. Self-organization: To involve the self-organization phenomenon in the process of cell maturation (component #8), cells secrete some signaling molecules, with concentration *C*_*s*_, which affects the propensities in the Gillespie algorithm and, consequently, the production of proteins. The signaling molecules diffuse in the medium according to the following reaction-diffusion equation:

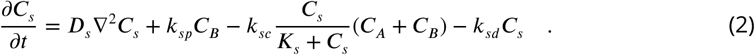

Here, *k*_*sp*_, *k*_*sc*_ and *k*_*sd*_ represent, respectively, the rate of production, consumption and decay of the signaling molecules and *D*_*s*_ is the diffusion coefficient of the signaling molecules. *C*_*A*_ and *C* _*B*_ respectively show the number of cells with phenotype ***A*** and ***B*** at each point of the medium. In our simulations, we used *D*_*s*_ = 10^−11^*m*^2^/*s, k*_*sp*_ = 0.01*kg*^−1^*s*^−1^, *k*_*sc*_ = 0.0001*kg*^−1^*s*^−1^, *k*_*sd*_ = 0.01*s*^−1^, and *K*_*s*_ = 0.01*m*^−3^.

In these simulations, the secreting cells are those with phenotype ***B***; hence, the production of signaling molecules is proportional to the amount of B cells. Since both phenotypes consume these molecules, the consumption depends on the number of both A and B cells. When ***B*** cells emerge, they secret signaling molecules, which diffuse in their environment. The minimum effective concentration of the signaling molecules at any location determines if a cellat the locationis affected by the signal, which would decrease the production of protein ***X*** and augment the production of protein ***Y***. Consequently, their surrounding cells would have less chance of producing protein ***X*** and their offspring is less likely to be in the domain of attraction of protein ***X***.

### Code availability

The software used to run all simulations was Matlab 2016 and the scripts are available at https://github.com/hasafdari/Noise_Driven_Cell_Differentiation (doi: https://doi.org/10.5281/zenodo.1227287).

## Acknowledgments

We had helpful discussions with Hamid Pezeshk and Amir Malekpour. Elahe Elahi, Steffen Rulands, Ricardo B. R. Azevedo, and Gábor Balázsi provided useful comments on the manuscript.

## Author contribution

MS designed research; RT and BG contributed to the initial idea; AK wrote the manuscript; HS and CP contributed to the methods section; MS and AK contributed to the introduction and the discussion; HS analyzed the data; HS and AK visualized the results. All authors read and approved the final manuscript.

## Additional information

The authors declare no competing interests.

## Appendix 1

### Movies

**Movie S1:** The change in the distribution of TFs within cells just before they they divide. Parameters used are the same as Fig 1.

Movie S2 and Movie S3 show 3-dimensional simulations of a community of cells in a layer. Simulation performed in a *L* × *L* × *h* cube and starts with one cell at the centre. The cells grow in volume; after reaching a critical volume they divide and the same as two dimensional case, their cytoplasmic content distributes between the two daughter cells.

**Movie S2:** The emergence of heterogeneity in the population of cells as a result of the presence of noise in the process of cell growth and division. The average amount of TFs in each cell at steady state is 25 The simulation started by one cell and continues over 13 generations, *L* = 130*μm* and *h* = 1.33*μm*. Since there is a single layer of cells, ***h*** corresponds to the diameter of a single cell.

**Movie S3:** The formation of a spatial organization as a result of the secretion of signaling molecules, which diffuse in their environment and affect the differentiation of the cells. The average amount of TFs in each cell at steady state = 25. The simulation started by one cell and continues over 13 generations, *L* = 130*μm* and *h* = 1.33*μm.*

**Appendix 1 Figure 1.**
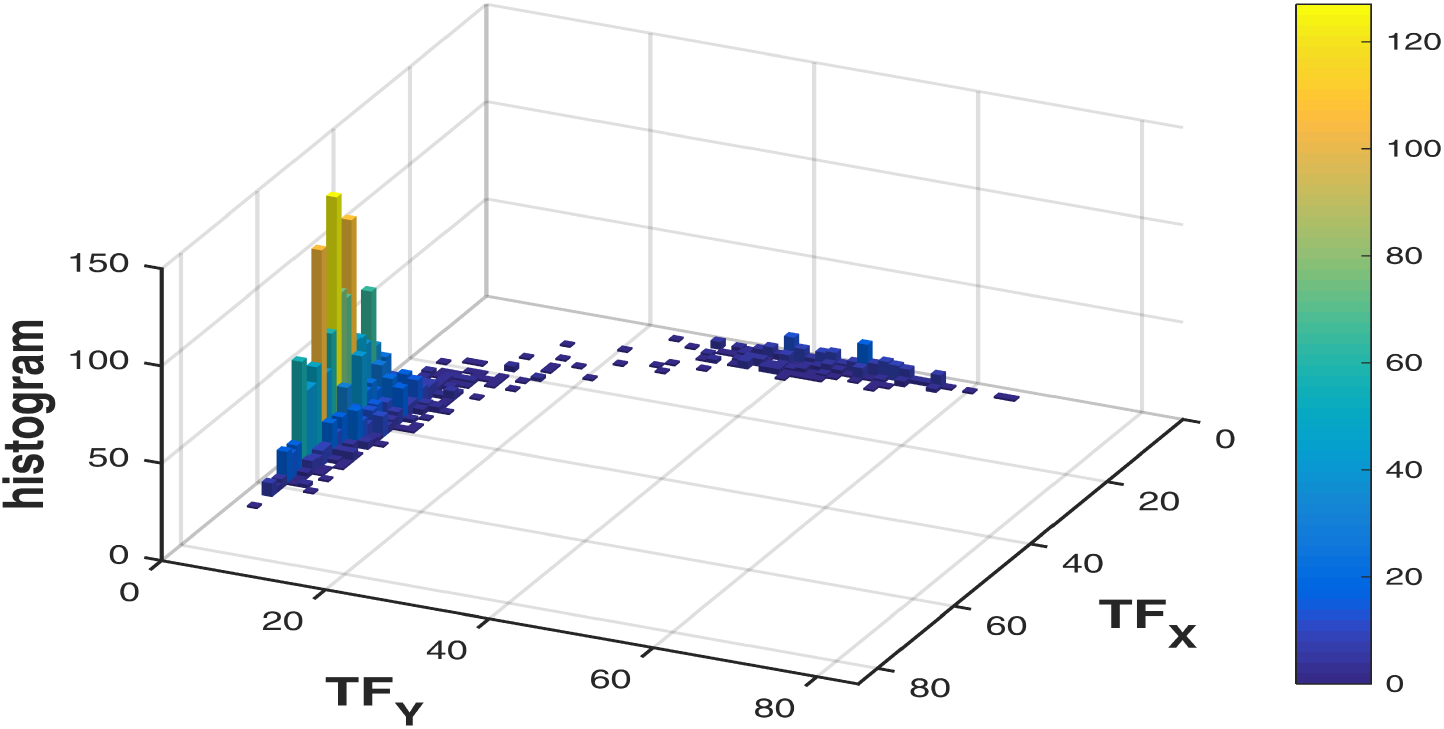
The final frame of Movie S1

**Appendix 1 Figure 2.**
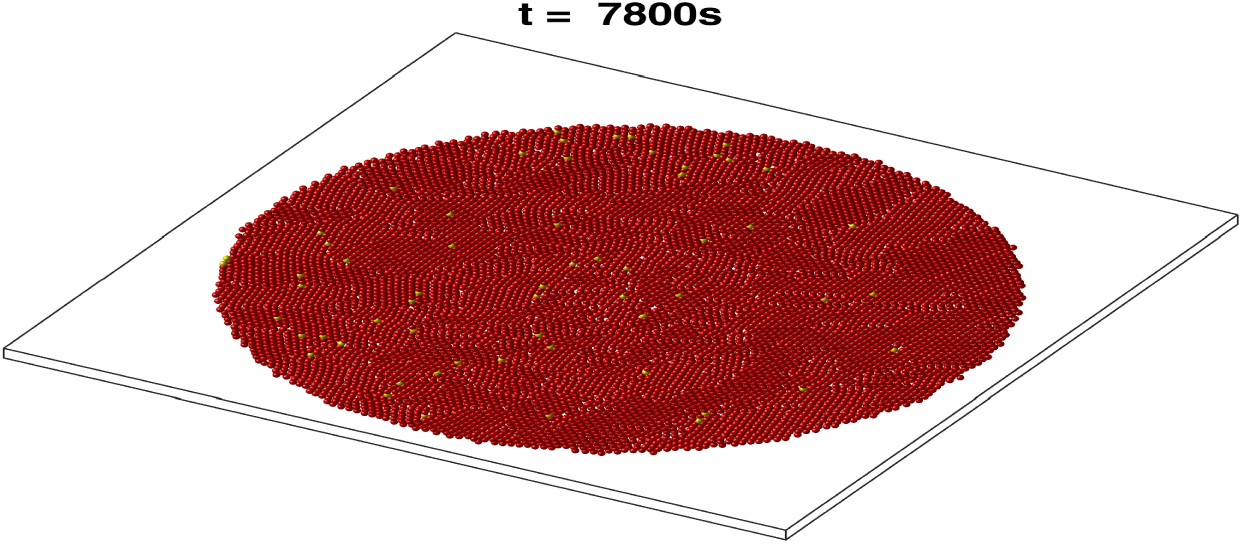
The final frame of Movie S2

**Appendix 1 Figure 3.**
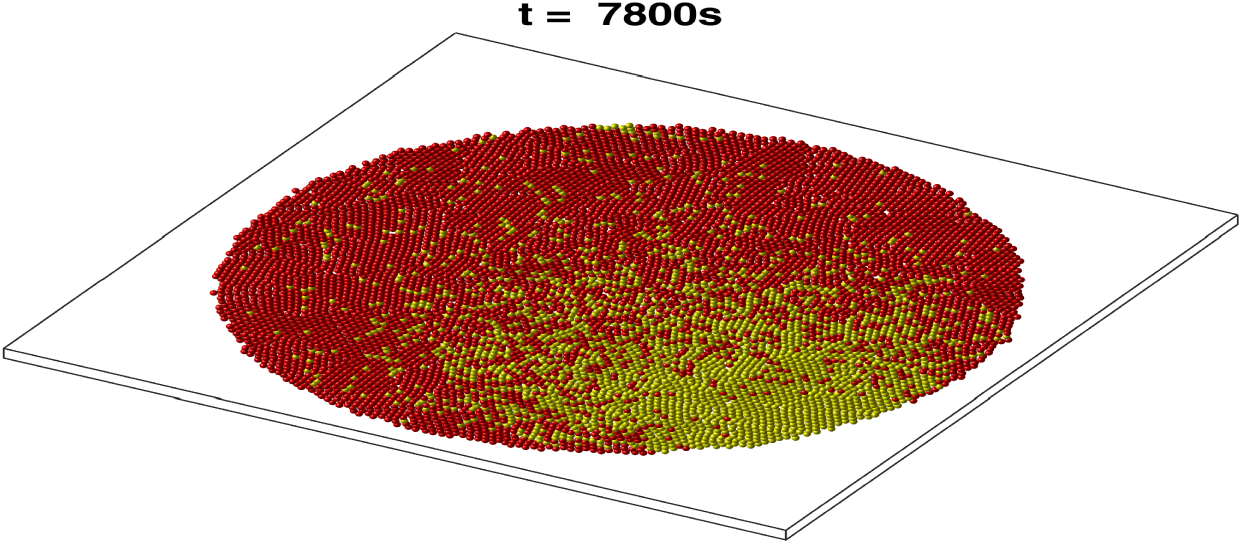
The final frame of Movie S3

## References

Allen B, Gore J, Nowak M. Spatial dilemmas of diffusible public goods. eLife. 2013; 2(0):e01169. doi: 10.7554/eLife.01169.

Axelrod K, Sanchez A, Gore J. Phenotypic states become increasingly sensitive to perturbations near a bifurcation in a synthetic gene network. eLife. 2015; 4:e07935. doi: 10.7554/eLife.07935.

Balázsi G, van Oudenaarden A, Collins JJ. Cellular Decision Making and Biological Noise: From Microbes to Mammals. Cell. 2011; 144(6):910–925. doi: 10.1016/j.cell.2011.01.030.

Betschinger J, Knoblich JA. Dare to Be Different: Asymmetric Cell Division in *Drosophila, C.elegans* and Vertebrates. Curr Biol. 2017 2017/06/07; 14(16):R674–R685. doi: 10.1016/j.cub.2004.08.017.

Bradshaw N, Losick R. Asymmetric division triggers cell-specific gene expression through coupled capture and stabilization of a phosphatase. eLife. 2015; 4:e08145. doi: 10.7554/eLife.08145.

Bumke MA, Neri D, Elia G. Modulation of gene expression by extracellular pH variations in human fibroblasts: A transcriptomic and proteomic study. Proteomics. 2003; 3(5):675–688. doi: 10.1002/pmic.200300395.

Carson E, Cobelli C. Modelling Methodology for Physiology and Medicine. Academic Press; 2000.

Chalancon G, Ravarani C, Balaji S, Alfonso M, Aravind L, Jothi R, Babu M. Interplay between gene expression noise and regulatory network architecture. Trends Genet. 2012; 28(5):221–232. doi: 10.1016/j.tig.2012.01.006.

Chang HH, Hemberg M, Barahona M, Ingber DE, Huang S. Transcriptome-wide noise controls lineage choice in mammalian progenitor cells. Nature. 2008 05; 453(7194):544–547.

Clevers H. Stem cells, asymmetric division and cancer. Nat Genet. 2005 10; 37(10):1027–1028.

Cortes MG, Trinh JT, Zeng L, Balázsi G. Late-Arriving Signals Contribute Less to Cell-Fate Decisions. Biophys J. 2018 2018/04/13; 113(9):2110–2120. http://dx.doi.org/10.1016/j.bpj.2017.09.012, xdoi: 10.1016/j.bpj.2017.09.012.

Dennett DC. Intuition Pumps And Other Tools for Thinking. W. W. Norton & Company; 2014.

Donlan RM. Biofilms: Microbial Life on Surfaces. Emerg Infect Diseases. 2002; 8(9):881 – 889.

Elowitz MB, Levine AJ, Siggia ED, Swain PS. Stochastic Gene Expression in a Single Cell. Science. 2002 08; 297(5584):1183.

Elowitz MB, Stanislas L. A synthetic oscillatory network of transcriptional regulators. Nature. 2000 01; 403(1038):1183.

Fisher RA. The Genetical Theory of Natural Selection. Oxford University Press; 1930.

Gardner A, Kalinka AT. Recombination and the evolution of mutational robustness. J Theor Biol. 2006; 241(4):707–715.

Gardner TS, Cantor CR, Collins JJ. Construction of a genetic toggle switch in *Escherichia coli*. Nature. 2000 01; 403(1038):1183.

van Gestel J, Vlamakis H, Kolter R. Division of Labor in Biofilms: the Ecology of Cell Differentiation. Microbiol Spectr. 2015; 3(2).

Ghaffarizadeh A, Flann N, Podgorski G. Multistable switches and their role in cellular Differentiation networks. BMC Bioinformatics. 2014; 15(S7):1–13. doi: 10.1186/1471-2105-15-S7-S7.

Gillespie DT. Exact stochastic simulation of coupled chemical reactions. J Phys Chem. 1977 12; 81(25):2340–2361. doi: 10.1021/j100540a008.

Hooshangi S, Bentley WE. LsrR Quorum Sensing “Switch” Is Revealed by a Bottom-Up Approach. PLOS Comput Biol. 2011 09; 7(9):1–11. https://doi.org/10.1371/journal.pcbi.1002172, xdoi: 10.1371/journal.pcbi.1002172.

Huang S. Non-genetic heterogeneity of cells in development: more than just noise. Development. 2009; 136(23):3853–3862. doi: 10.1242/dev.035139.

Huh D, Paulsson J. Non-genetic heterogeneity from stochastic partitioning at cell division. Nat Genet. 2010; 43(2):95–100. doi: 10.1038/ng.729.

Huh D, Paulsson J. Random partitioning of molecules at cell division. PNAS. 2011; 108(36):15004–15009. doi: 10.1073/pnas.1013171108.

Jan YN, Jan LY. Asymmetric cell division. Nature. 1998 04; 392:775 EP –. http://dx.doi.org/10.1038/33854.

Kepler TB, Elston TC. Stochasticity in Transcriptional Regulation: Origins, Consequences, and Mathematical Representations. Biophys J. 2001; 81(6):3116–3136. doi: http://dx.doi.org/10.1016/S0006-3495(01)75949-8.

Kimura M. On the evolutionary adjustment of spontaneous mutation rates. Genet Res. 1967; 9(1):23–34. doi: 10.1017/S0016672300010284.

Kirk DL. Volvox: molecular genetic origins of multicellularity and cellular Differentiation. New York: Cambridge University Press; 1998.

Kittisopikul M, Süel GM. Biological role of noise encoded in a genetic network motif. PNAS. 2010 07; 107(30):13300–13305.

Kreft JU, Booth G, Wimpenny JWT. BacSim, a simulator for individual-based modelling of bacterial colony growth. Microbiology. 1998; 144(12):3275–3287.

Kreft JU, Picioreanu C, Wimpenny JWT, van Loosdrecht MCM. Individual-based modelling of Biofilms. Microbiology. 2001; 147(11):2897–2912.

Kupiec JJ. A Darwinian theory for the origin of cellular Differentiation. Mol Gen Genet. 1997; 255(2):201–208. https://doi.org/10.1007/s004380050490, xdoi: 10.1007/s004380050490.

Lee C, Brangwynne C, Gharakhani J, Hyman A, Jülicher F. Spatial Organization of the Cell Cytoplasm by Position-Dependent Phase Separation. Phys Rev Lett. 2013; 111(8). doi: 10.1103/PhysRevLett.111.088101.

Lo M, Bulach DM, Powell DR, Haake DA, Matsunaga J, Paustian ML, Zuerner RL, Adler B. Effects of Temperature on Gene Expression Patterns in Leptospira interrogans Serovar Lai as Assessed by Whole-Genome Microarrays. Infect Immun. 2006 10; 74(10):5848–5859. doi: 10.1128/IAI.00755-06.

Losick R, Desplan C. Stochasticity and cell fate. Science. 2008; 320(5872):65–68.

Lynch M, Ackerman MS, Gout JF, Long H, Sung W, Thomas WK, Foster PL. Genetic drift, selection and the evolution of the mutation rate. Nat Rev Genet. 2016 11; 17(11):704–714.

Maamar H, Raj A, Dubnau D. Noise in Gene Expression Determines Cell Fate in *Bacillus subtilis*. Science. 2007; 317(5837):526–529. doi: 10.1126/science.1140818.

of Malmesbury TH. Leviathan or The Matter, Forme and Power of a Common Wealth Ecclesiasticall and Civil. London: Andrew Crooke; 1651.

Margolin W. Themes and variations in prokaryotic cell division. FEMS Microbiol Rev. 2000;.

Maynard Smith J, Szathmáry E. The Major Transition in Evolution. New York: Oxford University Press; 1995.

McDonald JH, Kreitman M. Adaptive protein evolution at the Adh locus in *Drosophila*. Nature. 1991 06; 351(6328):652–654.

Monahan L, Liew A, Bottomley A, Harry E. Division site positioning in bacteria: one size does not fit all. Front Microbiol. 2014; 5:19. doi: 10.3389/fmicb.2014.00019.

Morrison SJ, Kimble J. Asymmetric and symmetric stem-cell divisions in development and cancer. Nature. 2006 06; 441(7097):1068–1074.

Moussy A, Cosette J, Parmentier R, da Silva C, Corre G, Richard A, Gandrillon O, Stockholm D, Páldi A. Integrated time-lapse and single-cell transcription studies highlight the variable and dynamic nature of human hematopoietic cell fate commitment. PLOS Biology. 2017 07; 15(7):e2001867–. https://doi.org/10.1371/journal.pbio.2001867.

Mustonen M, Haimi J, Kesäniemi J, Högmander H, Knott KE. Variation in gene expression within clones of the earthworm Dendrobaena octaedra. PLOS ONE. 2017 04; 12(4):1–16. doi: 10.1371/journal.pone.0174960.

Nadell CD, Xavier JB, Foster KR. The sociobiology of Biofilms. FEMS Microbiol Rev. 2009; 33(1):206–24. doi: 10.1111/j.1574-6976.2008.00150.x.

Nanjundiah V. Cellular Slime Mold Development as a Paradigm for the Transition from Unicellular to Multicellular Life. In: Niklas KJ, Newman SA, editors. Multicellularity: Origins and Evolution Cambridge, Massachusetts: The MIT Press; 2016.p. 105–130.

Neildez-Nguyen TMA, Parisot A, Vignal C, Rameau P, Stockholm D, Picot J, Allo V, Le Bec C, Laplace C, Paldi A. Epigenetic gene expression noise and phenotypic diversification of clonal cell populations. Differentiation. 2008; 76(1):33–40. http://www.sciencedirect.com/science/article/pii/S0301468109600505, doi: https://doi.org/10.1111/j.1432-0436.2007.00219.x.

Novick A, Weiner M. Enzyme induction as an all-or-none phenomenon. PNAS. 1957 07; 43(7):553–566. http://www.ncbi.nlm.nih.gov/pmc/articles/PMC528498/.

Ostrowski EA, Shaulsky G. Learning to get along despite struggling to get by. Genome Biol. 2009; 10(5):218. doi: 10.1186/gb-2009-10-5-218.

Ozbudak EM, Thattai M, Kurtser I, Grossman AD, van Oudenaarden A. Regulation of noise in the expression of a single gene. Nat Genet. 2002 05; 31(1):69–73.

Paldi A. Stochastic gene expression during cell Differentiation: order from disorder? Cell Mol Life Sci. 2003; 60(9):1775–1778. https://doi.org/10.1007/s00018-003-23147-z, doi: 10.1007/s00018-003-23147-z.

Paliwal S, Iglesias PA, Campbell K, Hilioti Z, Groisman A, Levchenko A. MAPK-mediated bimodal gene expression and adaptive gradient sensing in yeast. Nature. 2007 03; 446(7131):46–51.

Perez-Carrasco R, Guerrero P, Briscoe J, Page KM. Intrinsic Noise Profoundly Alters the Dynamics and Steady State of Morphogen-Controlled Bistable Genetic Switches. PLOS Comput Biol. 2016 10; 12(10):1–23. doi: 10.1371/journal.pcbi.1005154.

Picioreanu C, Kreft JU, Van Loosdrecht MCM. Particle-based multidimensional multispecies biofilm model. Appl Environ Microbiol. 2004 May; 70(5):3024–3040.

Pickett-Heaps JD, Gunning BE, Brown RC, Lemmon BE, Cleary AL. The cytoplast concept in dividing plant cells: cytoplasmic domains and the evolution of spatially organized cell division. Am J Bot. 1999;.

Ptashne M. A Genetic Switch: Phage Lambda Revisited. Cold Spring Harbor Laboratory Press; 2004.

Queller DC, Strassmann JE. Beyond society: the evolution of organismality. Philos Trans R Soc Lond B Biol Sci. 2009 10; 364(1533):3143.

Rudel D, Sommer RJ. The evolution of developmental mechanisms. Dev Biol. 2003; 264(1):15–37. doi: http://dx.doi.org/10.1016/S0012-1606(03)00353-1.

Sanchez A, Golding I. Genetic determinants and cellular constraints in noisy gene expression. Science. 2013 12; 342(6163):1188–1193. doi: 10.1126/science.1242975.

Schluter J, Schoech AP, Foster KR, Mitri S. The Evolution of Quorum Sensing as a Mechanism to Infer Kinship. PLOS Comput Biol. 2016 04; 12(4):1–18. https://doi.org/10.1371/journal.pcbi.1004848, doi: 10.1371/journal.pcbi.1004848.

Sharifi-Zarchi A, Totonchi M, Khaloughi K, Karamzadeh R, Araúzo-Bravo MJ, Baharvand H, Tusserkani R, Pezeshk H, Chitsaz H, Sadeghi M. Increased robustness of early embryogenesis through collective decisionmaking by key transcription factors. BMC Syst Biol. 2015; 9(1):23. doi: 10.1186/s12918-015-0169-8.

Shea MA, Ackers GK. The OR control system of bacteriophage lambda. Journal of Molecular Biology. 1985; 181(2):211 – 230. doi: http://dx.doi.org/10.1016/0022-2836(85)90086-5.

Suzuki N, Furusawa C, Kaneko K. Oscillatory Protein Expression Dynamics Endows Stem Cells with Robust Differentiation Potential. PLOS ONE. 2011 11; 6(11):e27232–.

Vogt G. Stochastic developmental variation, an epigenetic source of phenotypic diversity with farreaching biological consequences. J Biosci. 2015 Mar; 40(1):159–204.

Wolk PC. Heterocyst formation. Annu Rev Genet. 1996; 30:59–78.

Wolpert L. Positional information and patterning revisited. J Theor Biol. 2011; 269(1):359–365. doi: http://dx.doi.org/10.1016/j.jtbi.2010.10.034.

Wu J, Tzanakakis E. Contribution of Stochastic Partitioning at Human Embryonic Stem Cell Division to NANOG Heterogeneity. PLOS ONE. 2012; 7(11):e50715. doi: 10.1371/journal.pone.0050715.

Xavier JB, Picioreanu C, van Loosdrecht MCM. A framework for multidimensional modelling of activity and structure of multispecies Biofilms. Environ Microbiol. 2005 Aug; 7(8):1085–1103. doi: 10.1111/j.1462-2920.2005.00787.x.

Xavier JdB, Picioreanu C, van Loosdrecht MCM. A general description of detachment for multidimensional modelling of Biofilms. Biotechnol Bioeng. 2005 Sep; 91(6):651–669. doi: 10.1002/bit.20544.

Yamagishi JF, Saito N, Kaneko K. Symbiotic Cell Differentiation and Cooperative Growth in Multicellular Aggregates. PLOS Comput Biol. 2016 10; 12(10):1–17. https://doi.org/10.1371/journal.pcbi.1005042, doi: 10.1371/journal.pcbi.1005042.

